# Quantitative Genomic Dissection of Soybean Yield Components

**DOI:** 10.1101/784538

**Authors:** Alencar Xavier, Katy M Rainey

## Abstract

Soybean is a crop of major economic importance with low rates of genetic gains for grain yield compared to other field crops. A deeper understanding of the genetic architecture of yield components may enable better ways to tackle the breeding challenges. Key yield components include the total number of pods, nodes and the ratio pods per node. We evaluated the SoyNAM population, containing approximately 5600 lines from 40 biparental families that share a common parent, in 6 environments distributed across 3 years. The study indicates that the yield components under evaluation have low heritability, a reasonable amount of epistatic control, and partially oligogenic architecture: 18 quantitative trait loci were identified across the three yield components using multi-approach signal detection. Genetic correlation between yield and yield components was highly variable from family-to-family, ranging from −0.2 to 0.5. The genotype-by-environment correlation of yield components ranged from −0.1 to 0.4 within families. The number of pods can be utilized for indirect selection of yield. The selection of soybean for enhanced yield components can be successfully performed via genomic prediction, but the challenging data collections necessary to recalibrate models over time makes the introgression of QTLs a potentially more feasible breeding strategy. The genomic prediction of yield components was relatively accurate across families, but less accurate predictions were obtained from within-family predictions and predicting families not observed included in the calibration set.

## Introduction

Soybean is a field crop of major importance due to its seed composition, containing approximately 40% protein and 20% oil. Soybean has restricted genetic basis (Mikel et al. 2010) and the rate of genetic improvement of soybeans has been reported to be 29 kg/ha/yr in North America (Rinker et al. 2014) and better breeding strategies are needed (Specht et al. 2014) to for soybean to reach its full genetic potential (Specht et al. 1999), especially considering its narrow genetic basis (Carter et al. 2004). A possible approach to increase grain yield is through trait dissection, breaking down yield into yield components. For instance, whereas modern cultivars have around 30 pods per plant (Kahlon et al. 2011), some accessions reach 200 pods per plant (Zhang et al. 2015).

Kahlon and Board (2012) contrasted cultivars released over the past few decades and observed that grain yield increases may have been triggered by changes in yield components the over time, particularly in pods and nodes. The number of pods and number of nodes are key yield-drivers (Robinson et al. 2009) that reflect the efficiency of complex physiological process (Board and Tan 1995). These yield components can be increased at the farm level with good agronomic practices and elite genetics (Board and Kahlon 2011, Kahlon et al. 2011). However, the labor-intensive phenotyping of counting soybean pods and nodes can restrict the number of entries and most studies have been conducted with a small number of genotypes (Egli and Bruening 2006, Robinson et al. 2009, Kahlon et al. 2011, Nico et al. 2019).

The first large-scale genetic assessment of complex traits was performed in the SoyNAM population, where 5600 genotypes from 40 bi-parental families sharing a common parent were phenotyped for various agronomic traits (Xavier et al. 2016, Xavier et al. 2017a, Diers et al. 2018). Whereas soybeans have constrained genetic diversity (Carter et al. 2004), the SoyNAM is a relatively rich panel of locally adapted genotypes that represents an invaluable resource for the breeding community. From a preliminary analysis in the SoyNAM population, Xavier et al. (2017a) indicated that grain yield can be genetically improving yield components, rapid canopy development, and by increasing the length of the reproductive period. The latter is a function of days to flowering and days to maturity, both traits controlled by a few major genes (Watanabe et al. 2009, 2011, Xia et al. 2012, Langewisch et al. 2014). The genetic architecture of canopy development was recently described by Xavier et al. (2017b) and Kaler et al. (2018). However, the in-depth genetic architecture of yield components had not been characterized with sufficient power and resolutions.

This study aims to conduct a set of quantitative genetic analyses performed with genome-wide markers to decipher the underlying architecture of yield components and assess potential breeding applications. Our evaluation approach includes comparing different strategies for genomic prediction within and across family; perform genomic covariance analysis to uncover the pleiotropy between yield and yield components, as well as the amount of genetic variation attributed to epistasis and genotype-by-environment interactions; and multi-approach association studies to identify regions containing QTLs with the potential to be deployed for marker-assisted selection.

## Methods

### Population

The panel under evaluation is a nested association panel, namely the SoyNAM populations, where the standard parent IA3023 (Dairyland DSR365 x Pioneer P9381) was crossed to 40 founder parents that attempt to capture the diversity of public germplasm, each family comprising approximately 140 individuals. Among the 40 founder parents, 17 lines are U.S. elite public germplasm, 15 have diverse ancestry, and eight are planted introductions.

The descriptions of parents are available https://www.soybase.org/SoyNAM/. The population’s maturity ranged from late maturity group II to early maturity group IV. More details about the population composition are available in Diers et al. (2018) and Xavier et al. (2018). After quality control based on segregation patterns, 5363 individuals were used for this study.

### Experimental design

The experiment was conducted under a modified augmented design, with a 7:1 lines-to-check ratio, in two Purdue research centers: Throckmorton-Purdue Agricultural Center (TPAC) located in Throckmorton, Indiana, and at the Agronomy Center for Research and Education (ACRE) in West Lafayette, Indiana. The experiments were planted during the third week of May in two-row plots (2.9m × 0.76m), at a density of approximately 36 plants m^-2^. The phenotypes were collected in 10 local environments, these being distributed as 4 fields in 2013, 4 fields in 2014 and 2 fields in 2015. Experimental units were allocated into 6 fields allocated in three years (2013-2015). In 2013 and 2014 the experiments were conducted at the ACRE farm, where each local environment contained all 40 families with 35 recombinant inbred lines (RIL) per family, that is, one-quarter of the total number of RILs. The same checks were used across fields. In 2015, the experiments were conducted on 6 of the 40 SoyNAM families in two locations, ACRE and TPAC, with two replicates per location.

### Phenotyping

The number of pods and nodes was counted in the main stem, between phenological stages R5 and R7, averaging the counts of 3, 6 and 4 representative plants per plot in 2013, 2014 and 2015 respectively (Xavier et al. 2016). The number of pods per node was obtained by the ratio. Grain yield was collected at harvest, converting the grain weight from individual plots to bushels per acre adjusted to 13% grain moisture. The number of days to maturity (Fehr et al. 1971) was collected by scoring the plots every 3 days from the time where the first mature plot was observed, using back-and-forth scoring to assign the plots that matured between scoring dates.

### Genotyping

The genetic information was collected from Illumina SoyNAM BeadChip SNP array specially designed for SoyNAM, comprising 5305 SNP markers selected from the sequencing of all 41 parental lines (Song et al. 2017). Missing loci were imputed using a hidden Markov model and removed markers with minor allele frequency below 0.05 using the R package NAM (Xavier et al. 2015). A total of 4240 SNPs was used for genomic analysis.

### Genetic merit

The genetic values were estimated as the best linear unbiased predictors (BLUP), as a random term of a mixed model. The mixed linear model was fitted with variance components based on restricted maximum likelihood (REML), computed using the R package lme4 (Bates et al. 2014). The linear model used to model genetic values:

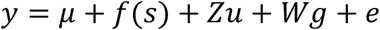

Where the response variable *y* was modeled as a function of an intercept *μ*, spatial covariate *f* (*s*) based on a moving-average of neighbor plots as described by Lado et al. (2013) implemented in the functions NNscr/NNcov of the R package NAM (Xavier et al. 2015), a random effect *Zu* to capture the genetic effects of individual lines, namely the genetic effects, assumed to be normally distributed as *u* ∼ *N*(0, *σ*^2^_*u*_), a nuisance random effect *Wg* to capture the local environment effects, as normally distributed as *g* ∼ *N*(0, *σ*^2^_*g*_), and a vector *e*of residuals, normally distributed *e* ∼ *N*(0, *σ*^2^_*e*_). The inverse phenotypic variance was computed for each environment and used as observation weights to account for the heteroscedasticity among trials. Although the checks were not explicitly included in the genetic merit model, these were invaluable for the spatial correction of the field plot variation. Broad-sense heritability (H) was estimated from the REML variance components as:

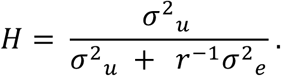

Where *r* is the average number of replicates per entry. The reliability of the *j*^*th*^ genotype (*H*_*j*_) was used to deregress (Garrick et al. 2009) its corresponding BLUP (*u*_*j*_) to obtain the genetic values in natural scale (*y*_*j*_ = *u*_*j*_/*H*_*j*_). This procedure to unshrink BLUPs precludes the downstream analyses to be performed upon a vector of phenotypes with heterogeneous degrees of shrinkage, which may lead to biased results. The narrow-sense heritability estimates (*h*^2^) were based on the following SNP-BLUP model:

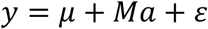

Where *y* correspond to the genetic values, modeled as a function of an intercept *μ*, matrix with SNP information and marker effects (*Ma*), and the vector of residuals (*ε*). Both marker effects and residuals were assumed to be normally distributed with variances *σ*^2^_a_ and *σ*^2^_*ε*_, respectively. The narrow-sense heritability was computed under two scenarios: 1) deploying all markers and 2) only with the markers found to be associated with yield components. The narrow-sense heritability was estimated as follows:

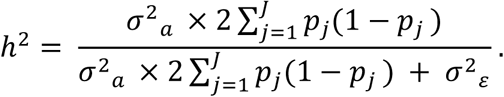

### Polygenic epistasis

We performed a within-family variance component analysis to determine the amount of variability jointly explained by additive and additive-by-additive epistasis. For that, we fit a kernel-based model referred to as the G2A model (Zeng et al. 2005). Variance components were estimated using REML estimates (Misztal 2008). The analysis followed the linear model:

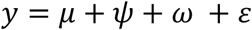

Where *y* correspond to the genetic values, modeled as a function of an intercept *μ*, additive genetic values, *ψ* ∼ *N*(0, *Kσ*^2^_*ψ*_), additive epistatic value, *ω* ∼ *N*(0, *Qσ*^2^_ω_), and the vector of residuals, *ε* ∼ *N*(0, *Iσ*^2^_*ε*_). The relationship matrices were built in accordance to Zeng et al. (2005) and Xu (2013). The additive genetic relationship matrix was obtained by the cross-product of the centralized marker matrix (*M*) with centralized trace, thus *K* = *MM*′*α* with *α* = *n* × *Tr*(*MM*′)^−1^, and the additive epistatic relationship matrix was computed by the additive Hadamard product with centralized trace, thus *Q* = (*MM*′#*MM*′)*α* with a normalizing factor *α* = *n* × *Tr*(*MM*′#*MM*′)^−1^.

### Multivariate analysis of pleiotropy and stability

Multivariate analysis, namely genetic and additive genetic correlations, allows exploration of the interaction between traits across years (pleiotropy) or within trait between years (stability or genotype-by-environment correlation). The genetic correlations within-family were obtained as the Pearson’s correlation between the BLUPs of yield and yield components for pleiotropy analysis, as well as the correlations of yield components from year to year for stability analysis. We estimated the additive genetic correlation between yield and yield components for pleiotropy analysis, and yield components across years for stability analysis, for each of the 40 families using a multivariate GBLUP model. The GBLUP model was fit with REML variance components. For the multivariate polygenic analysis, we fitted the following multi-trait model:

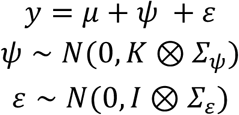

Where, under multivariate settings, *y* = {*y*_1_, *y*_2_, …, *y*_k_} correspond to the genetic merits, modeled as a function of their corresponding intercepts, *μ* = {*μ*_1_, *μ*_2_, …, *μ*_k_}, the additive genetic values, *ψ* = {*ψ*_1_, *ψ*_2_, …, *ψ*_k_}, and the residuals, *ε* = {*ε*_1_, *ε*_2_, …, *ε*_k_}. With respect to the model variances, *K*is the relationship matrix defined in the previous model, the additive covariances *Σ*_*ψ*_ is a dense *k* × *k* matrix where the *ij* cell corresponds to the additive genetic covariance *σ*_*ψ*_(*i, j*) between *i*^*th*^ and *j*^*th*^ traits, and the residual covariance was assumed to be diagonal *Σ*_*ε*_ = *diag*(*σ*^2^_*ε*1_, *σ*^2^_*ε*2_, …, *σ*^2^_*ε*k_). Additive genetic correlations were estimated from the covariance components as ρ_*ψ*_(*i, j*) = *σ*_*ψ*_(*i, j*)/[*σ*_*ψ*_(*i*)*σ*_*ψ*_(*j*)]. From the genetic correlations and heritabilities, the efficiency of indirect selection (Falconer and Mackay, 1996) using *i*^*th*^ trait to select the *j*^*th*^ trait was estimated as *E* = *h*_*j*_^−2^ *h*_*i*_ ^2^ *ρ*_*ψ*_(*i, j*).

### Association studies

Since various signal detection strategies may capture different QTLs (Yang et al. 2018), three complementary methodologies of genome-wide association studies were deployed in this study: Single marker analysis, implemented in the R package NAM (Xavier et al. 2015), whole-genome regression BayesCpi (Habier et al. 2011) implemented in the R package bWGR, and random forest implemented in the R package ranger (Wright and Ziegler 2015). A brief description of the methods is provided below.

#### 1) Mixed Linear Model (MLM)

This method of an association study is based on the likelihood ratio between a model containing the marker of interest (full model) and a model without the marker (reduced model). Both models include a polygenic term that accounts for the population structure. The statistical model that describes this association study is tailored to NAM populations (Xavier et al. 2015) and follows the linear model:

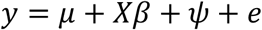

Where *y* correspond to the genetic values, modeled as a function of an intercept (*μ*), the matrix containing the interaction between the SNP information and family for the target marker under evaluation (*X*), the vector of marker effect within family *β* ∼ *N*(0, *σ*^2^_*β*_), the vector of independent residuals, *ε* ∼ *N*(0, *σ*^2^_*ε*_), and the polygenic term defined previously, *ψ* ∼ *N*(0, *Kσ*^2^_*ψ*_), which parametrizes the genetic covariance among individuals through the full-ranking genomic relationship matrix *K*. Bonferroni thresholds were utilized to account for multiple testing and mitigate false-positives, yielding a two-sided threshold of −log10(0.025/4240)=5.23. The association model was fit with REML variance components (Xavier et al. 2015).

#### 2) Whole-genome regression (WGR)

Designed primarily for prediction, WGR methods fit all markers are together at once. The prior distribution of marker effects follows a mixture of distributions to perform feature selection. The association statistics are based on the posterior probability of each marker to be included in the model, or “model frequency”. The model of choice, BayesCpi, assumes each marker has a probability *π* of being included in the model, where the parameter *π* is estimated in each MCMC iteration. Markers reached statistical significance if 1-*π* was smaller than a two-sided threshold of *α* = 0.05, which translates into a threshold for the Manhattan plot of −log10(0.025)=1.6. The linear model that describes BayesCpi is the following:

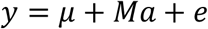

Where *y* correspond to the genetic values, modeled as a function of an intercept (*μ*), the matrix containing the all SNP information (*M*) and the vector of all marker effects jointly estimated (*a*), which followed a mixture of distributions, having probability *π* of having null effect and probability 1 − *π* or being normally distributed as *N*(0, *σ*^2^_*β*_), and the vector of independent residuals, *ε* ∼ *N*(0, *σ*^2^_*ε*_). The marker and residual variances were assumed to follow an inverse scaled chi-squared distribution, *σ*^2^_*β*_ ∼ *χ*^2^(*S*_*β*_, *v*_0_) and *σ*^2^_*ε*_ ∼ *χ*^2^(*S*_*ε*_, *v*_0_), assuming *v*_0_ = 5prior degrees of freedom and shape parameters computed assuming prior heritability of 0.5 (Pérez and de Los Campos, 2014), thus *S*_*β*_ = 0.5 *σ*^2^_*y*_ *MSx*^−1^(1 − *π*)^−1^and *S*_*ε*_ = 0.5 *σ*^2^_*y*_. The model was fit with 20000 MCMC iterations, discarding the initial 2000 iterations, and no thinning, such that the posterior means were computed by averaging 18000 MCMC iterations.

#### 3) Random forest regression (RFR)

Random forest is a non-parametric regression derived from the bootstrapping aggregation of decision trees built from subsets of data and parameters. The association statistics of RFR is based on feature importance (Botta et al., 2014). The forest was grown with 10000 decision trees. The trees were built having as starting point 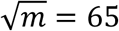 SNPs sampled at random with replacement. The metric of variable importance was the ‘impurity’ index, which is a measure of the out-of-bad explained variance. Because there is no objective way of defining an association threshold for significant SNPs, we estimated the global empirical threshold (Doerge and Churchill 1996) based on 1000 permutations (*α* = 0.05), thus making no assumptions about the distribution of the associations.

### Cross-validation studies

Cross-validations were performed for each yield component. Due to the known population structure of the SoyNAM, three types of cross-validations were performed: Within-family, across-family, and leave-family-out. Within- and across-family validations were performed as 5-fold cross-validation, randomly selecting 80% of the data as a calibration set and using the remaining 20% as a prediction target. The sampling and prediction procedure is repeated 25 times. Leave-family-out validation used 39 families to predict the family left out, and the procedure is performed to all 40 families. The prediction statistic is the predictive ability (PA), as the correlation between predicted and observed values.

The cross-validation was performed using the functions *emCV* of the R package bWGR (Xavier et al. 2017). In accordance with the genomic prediction benchmark proposed by Daetwyler et al. (2013), two statistical models evaluated in this study were GBLUP (VanRaden 2008) and BayesB (Meuwissen et al. 2001). The GBLUP model was fitted as a ridge regression with REML variance components, and the BayesB assumes that markers effects follow a mixture of distribution, where the *j*^*th*^ marker had probability *π* = 0.95 of having null effect and probability 1 − *π* of being normally distributed as *N*(0, *σ*^2^_*βj*_), variances were assumed to follow an inverse scaled chi-squared distribution, *σ*^2^_*βj*_ ∼ *χ*^2^(*S*_*β*_, *v*_0_)and *σ*^2^_*ε*_ ∼ *χ*^2^(*S*_*ε*_, *v*_0_), assuming *v*_0_ = 10 prior degrees of freedom and shape parameters computed as *S*_*β*_ = 0.5 *σ*^2^_*y*_ *MSx*^−1^and *S*_*ε*_ = 0.5 *σ*^2^_*y*_.

### Data availability

All phenotypic and genotypic data are available in the R package SoyNAM.

## Results

The SoyNAM provided reasonable variation for the three yield components. The phenotypic distributions of the yield components for each of the SoyNAM families is presented in Figure 1. The mean and standard deviation across families is provided in Table 1, alongside the broad- and narrow-sense heritability estimated across families.

**Table 1:**
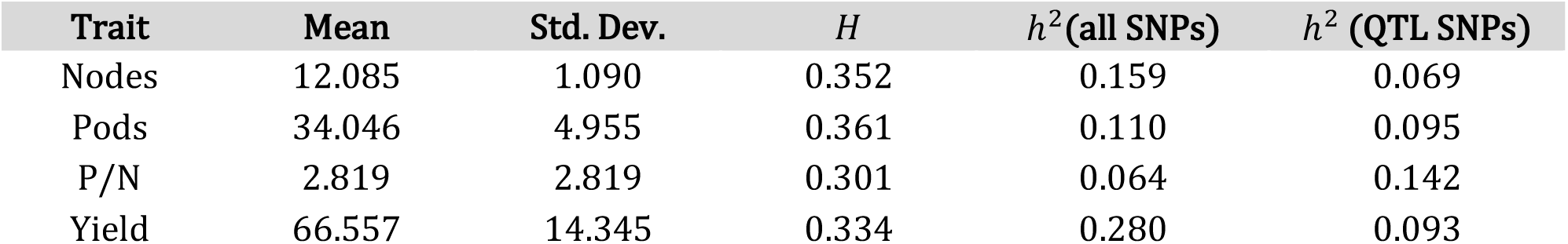
Trait distribution (mean and standard deviation) and genetic metrics: broad-sense heritability (*H*), narrow-sense heritability (*h*^2^) estimated using all SNPs and the subset of significant SNPs.

**Figure 1.**
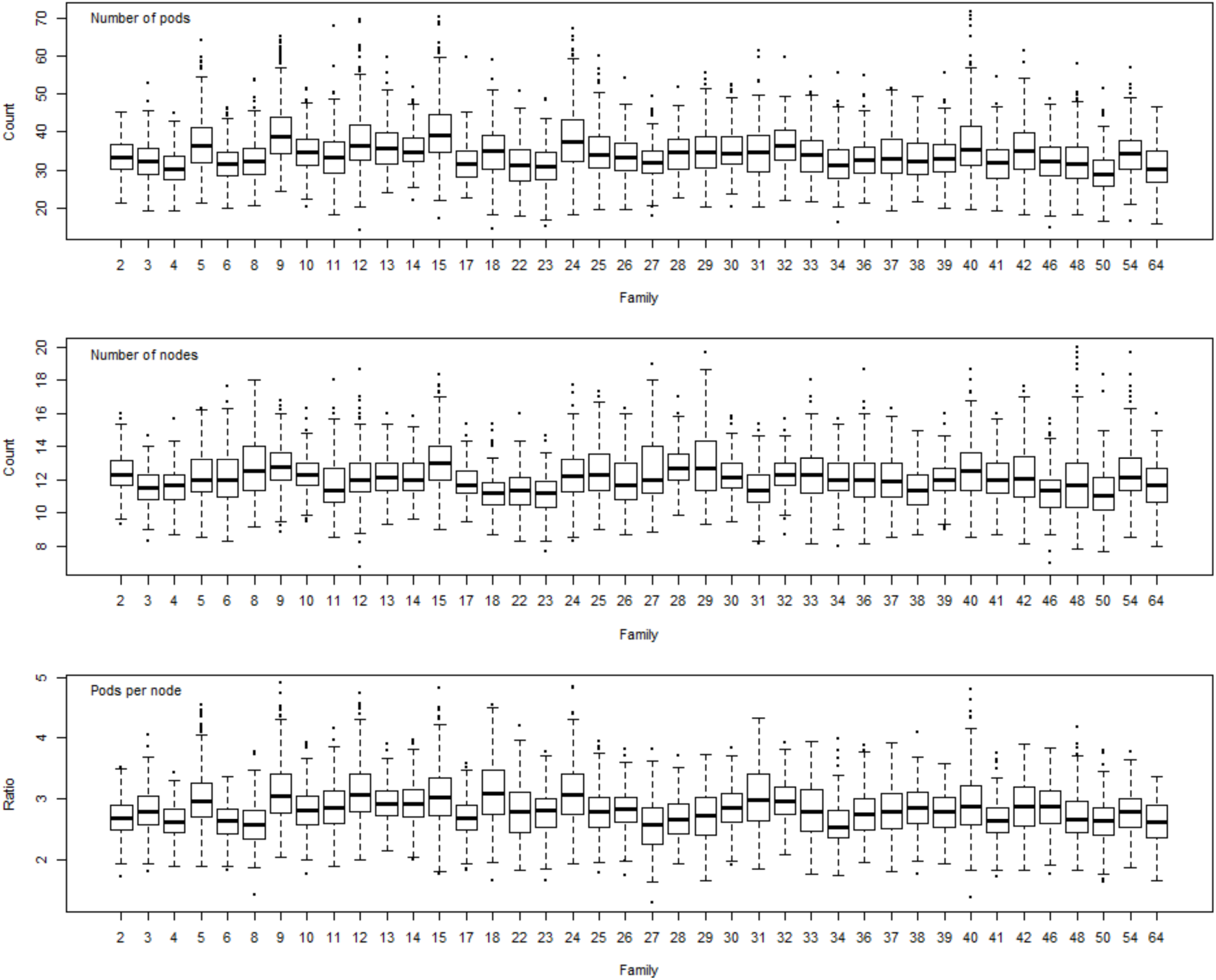
Phenotypic distribution of the pod number (top), node number (center) and pods per node (bottom).

The broad-sense heritability of the number of pods and nodes was slightly higher than the broad-sense heritability of yield, however, the narrow-sense heritability of yield was almost twice as large as the number of nodes, and almost three times higher than the narrow-sense heritability of the number of pods. The narrow-sense heritability estimated from the 18 markers found associated with yield components recovered almost entirely the narrow-sense heritability of the number of pods, but just a third of the heritability of the number of nodes and grain yield. And, surprisingly, the narrow-sense heritability of the ratio of pods per node was higher when only the significant markers were used.

### Association analysis

The genome-wide screening for segments associated to yield components is presented in Figure 2. Regions associated with the number of pods were located in chromosomes 3, 5, 14 and 19; significant associations for node number were observed in chromosomes 2, 3, 5, 6, 14, 18 and 19; and regions associated with pods per node were detected in chromosomes 3, 7, 12 and 19. The summary of the associated regions is presented in Table 2, alongside the impact of each significant marker on the yield components, grain yield and days to maturity. Except for the association between the marker Gm02_6396340 and the number of nodes, our study did not find any other consensus QTL detected by all three association methods for any of the variance components. All three yield components had significant associations in chromosomes 3 and 19, and the marker Gm19_1587494 was associated with all three traits. From the associated markers, Gm13_14346156 had the highest impact on grain yield, potentially increasing yield as much as 0.6 bushels per acre.

**Table 2:** Summary of association studies: SNP at the peak of each QTL; corresponding trait and method from which the QTL was identified, and the least squared effect of the SNP for each yield components, yield and days to maturity.

| SNP | GWAS (Fig. 2) | Pods | Nodes | P/N | Yield | Maturity |
| --- | --- | --- | --- | --- | --- | --- |
| Gm02_6396340 | B,D,E,F | -0.2607 | -0.4092 | -0.0297 | -0.4653 | -0.1617 |
| Gm03_2182974 | A,D,E | -0.2508 | -0.3828 | -0.0297 | -0.1815 | -0.3333 |
| Gm03_46533591 | A,B,G,H | -0.3168 | -0.1155 | -0.3399 | 0.0726 | 0.0198 |
| Gm05_914933 | B | -0.2013 | -0.1716 | -0.1023 | 0.1452 | 0.0198 |
| Gm05_3661638 | B,C | 0.1287 | 0.2475 | -0.0495 | -0.2277 | 0.3036 |
| Gm06_47199506 | D | 0.099 | 0.2112 | -0.0528 | 0.2211 | 0.0561 |
| Gm07_7868756 | G,H | -0.0198 | 0.1782 | -0.1848 | 0.0462 | -0.033 |
| Gm12_2838455 | I | 0.0693 | 0.1452 | -0.0363 | -0.0891 | 0.0792 |
| Gm13_14346156 | D | 0.1914 | 0.2607 | 0.0528 | 0.6171 | 0.0726 |
| Gm14_743883 | A | 0.2277 | 0.1584 | 0.1782 | 0.1815 | 0.1254 |
| Gm14_917668 | B | 0.2277 | 0.1914 | 0.1452 | 0.1056 | 0.2244 |
| Gm14_2322106 | D | 0.1683 | 0.2739 | -0.0099 | 0.3993 | 0.3168 |
| Gm15_5446785 | H | -0.0858 | 0.1386 | -0.2343 | -0.2376 | -0.0231 |
| Gm18_2357823 | D,E | -0.1122 | -0.2409 | 0.0495 | -0.2739 | -0.0165 |
| Gm18_57370051 | D,E | 0.2343 | 0.2772 | 0.0759 | 0.1386 | 0.2178 |
| Gm19_1496625 | B,E,I | -0.429 | -0.3795 | -0.2772 | -0.3366 | 0.0198 |
| Gm19_1587494 | B,C,F,H,I | -0.4257 | -0.3333 | -0.3102 | -0.1452 | 0.0891 |
| Gm19_1991181 | E | -0.363 | -0.3432 | -0.2112 | -0.1452 | 0.099 |

**Figure 2.**
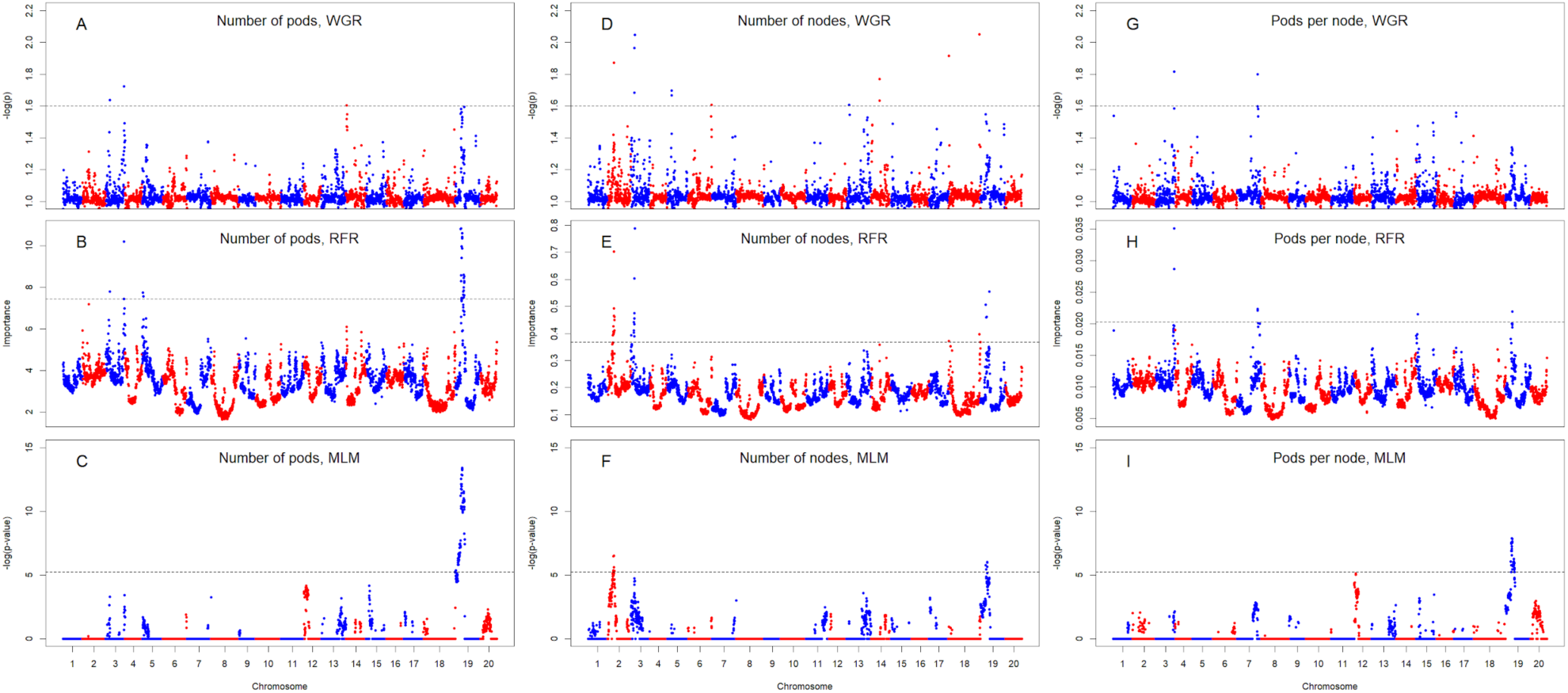
Genome-wide association studies of pod number (A,B,C), node number (D,E,F) and pods per node (G,H,I), performed through three methodologies: whole-genome regression (A,D,G), random forest regression (B,E,H), and mixed linear model (C,F,I).

### Prediction analysis

The outcome of the prediction analysis is presented in Figure 3. Predictions within-family provided lower correlations than leave-family-out, and across-family predictions provided the most predictive scenario. All three yield components had similar heritabilities (Table 1) and, consequently, similar prediction accuracies. For the different cross-validation scenarios, correlations around 0.05, 0.08 and 0.21 were observed for predictions within-family, leave-family-out, and across-families, respectively. BayesB provided a slightly higher predictive ability than GBLUP across cross-validation scenarios, providing an increase in predictability of as much as 0.02. However, the differences in predictive ability were negligible, in agreement with previous results (Xavier et al. 2016). The slightly advantageous performance of BayesB suggests that some QTLs contribute to the prediction of yield components, but a polygenic model captures most of the genomic signal.

**Figure 3.**
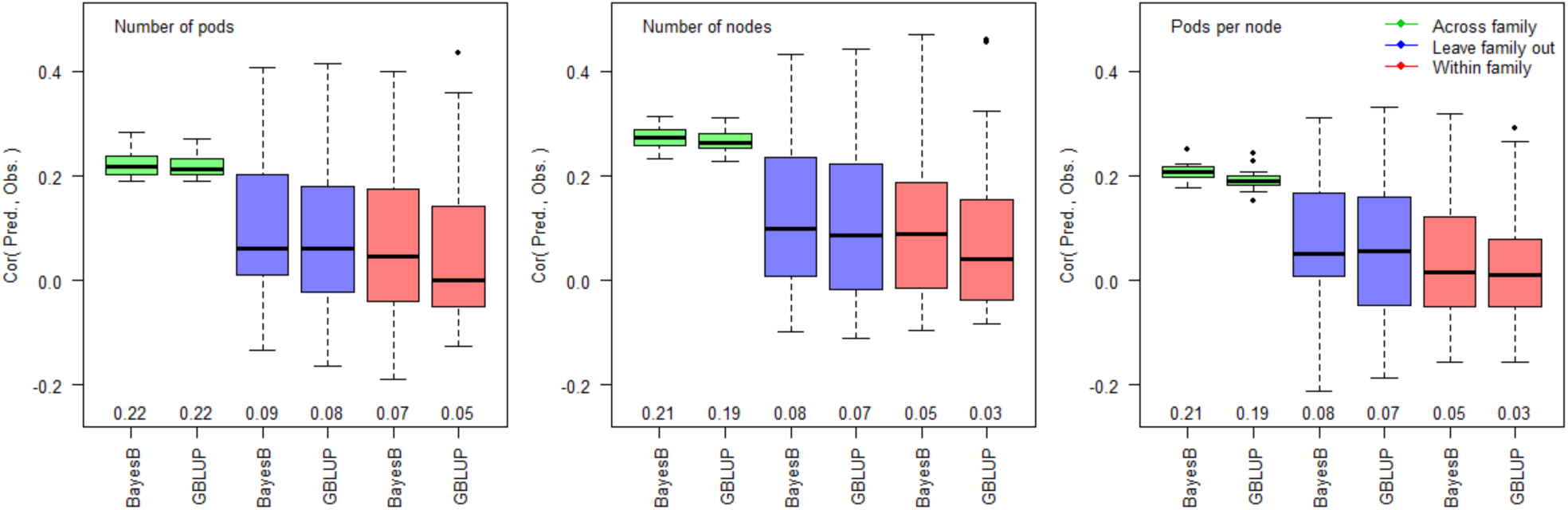
Boxplot of predictive ability of pod number (left), node number (center) and pods per node (right), where two prediction models (BayesB and GBLUP) tested three cross-validations strategies: across-family (green), leave-family-out (blue) and within-family (red).

### Variance decomposition

The proportion of variance explained by additivity and epistasis for individual families is presented in Figure 4. The additive fraction of the genetic variance computed using G2A kernels is comparable to the narrow-sense heritability estimated across families (Table 1). All three variance components presented similar average polygenic architecture, having the additive and epistatic components ranging from 0 to approximately 50%, but the estimates were highly variable from family to family. The additive component averaged 7.46%, 9.03%, and 6.18%; the epistatic component averaged 7.92%, 7.02%, and 7.77%, and the total genomic heritability (additive+epistatic components) averaged 15.38%, 16.05%, and 13.95% for the number of pods, nodes, and pods per node, respectively. Many families provided near-zero genetic control for yield components, in agreement with the low within-family predictive ability (Figure 3).

**Figure 4.**
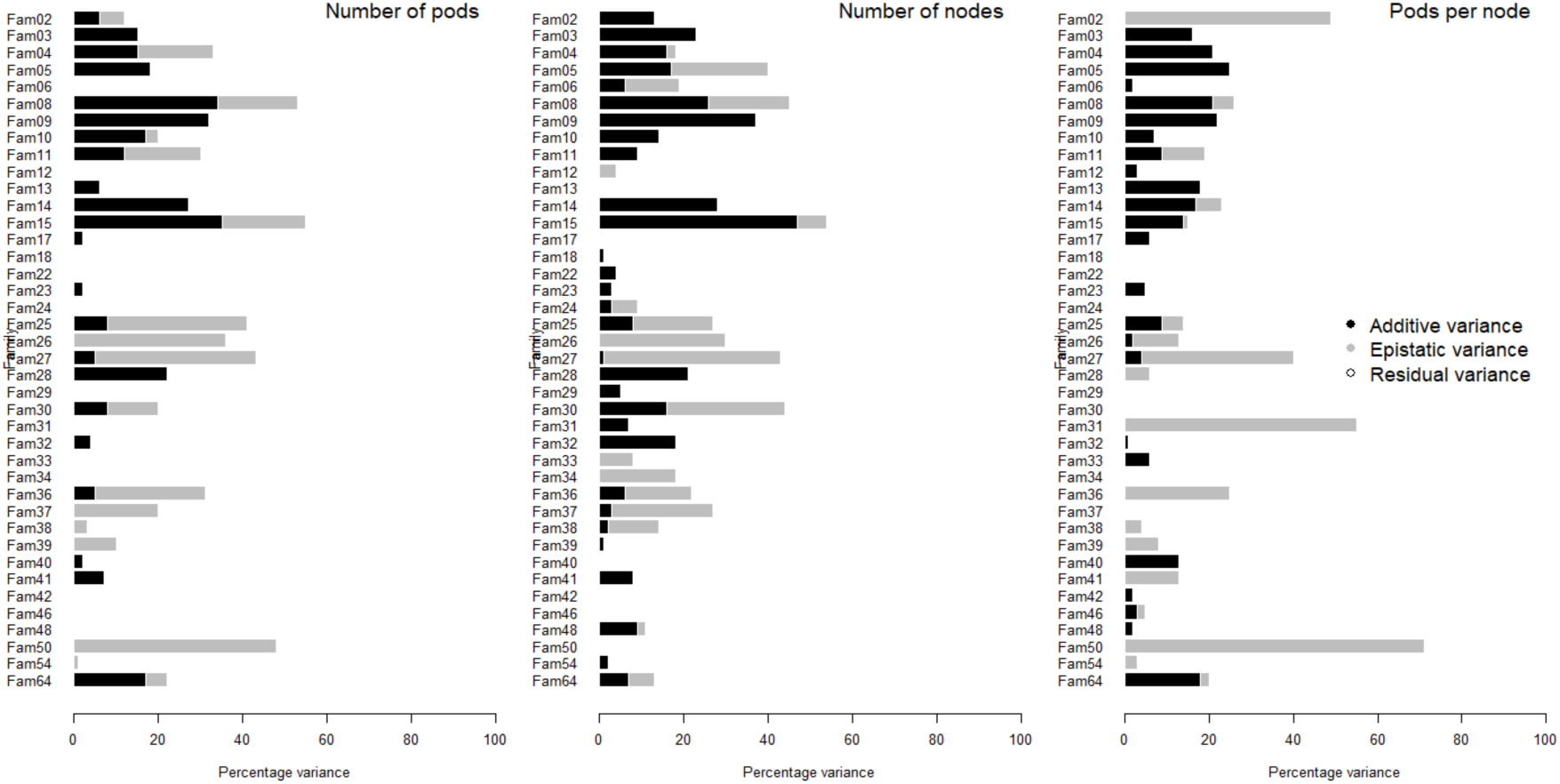
Barplot of the proportion of variance explained by different genetic components of pod number (left), node number (center) and pods per node (right) by family. Additive (black), epistatic (gray) and residual (white) variances.

### Genetic correlations and indirect selection

The within-family genetic and additive genetic correlations between yield components and yield, as well as yield components stability, are presented in Figure 5. Whereas the average correlations between yield components and yield are relatively small (Figure 5A), there is a large variation from family to family, which indicates that some families could benefit from the selection of yield components.

**Figure 5.**
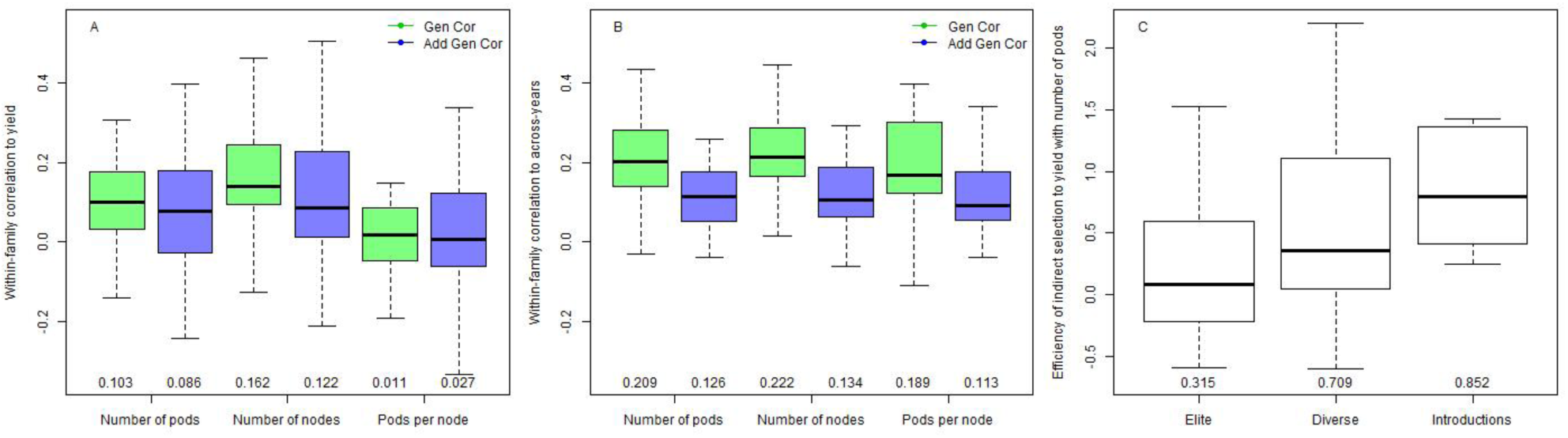
Boxplot displaying the dispersion of within-family genetic and additive genetic correlation between yield components and grain yield (A); the within-family genetic and additive genetic genotype-by-environment correlation (B); the efficiency of indirect selection to yield using pods, breakdown by germplasm background (C).

From the three yield components, the number of pods was the only trait with the efficiency of indirect selection that departed from zero (data not presented), so we broke down the efficiency of indirect selection based on pod counts by the genetic background of the SoyNAM founder (Figure 5C). Families with non-elite genetic backgrounds are more likely to benefit, and the indirect selection based on pods was more effective than on yield itself in 10 families (*E* > 1).

## Discussion

### Architecture overview

The number of pods and nodes, as the ratio pods per node, are key yield components in soybean (Herbert and Litchfield, 1982). Understanding how such traits work may provide insight into better strategies to increase yield and yield stability (Xavier et al. 2017a). In soybeans, these yield components have low heritability, both in the broad and narrow sense, and have partially oligogenic architecture, where the genomic control is jointly explained by a set of QTLs and polygenic terms (Figures 2 and 3). In addition, within-family analysis indicates that some populations display more epistatic than additive control under the polygenic model (Figure 4), whereas other families presented no genetic control whatsoever.

### QTLs

Successful mapping of markers associated with complex traits relies on the size and variability of the mapping population. Our study was conducted on the SoyNAM, a large population designed to optimize power and resolution. Yet, only a small number of QTLs were detected. Previous mapping studies on yield components have relied on non-experimental panels with a highly diverse genetic background. The studies of Hao et al. (2011), Hu et al. (2014), Zhang et al. (2015) and Fang et al. (2017) assessed 191, 113, 219 and 809 genotypes, respectively, including landraces and wild accessions.

Among the studies on diverse backgrounds, Fang et al. (2017) found a QTL for pod and node numbers in close proximity to our QTL peak on chromosome 06, marker Gm06_47199506. The pod number QTLs detected by Hu et al. (2014) were located in chromosomes 3, 5 and 6, in overlapping regions to signals Gm03_2382974, Gm03_46533591, Gm05_914933, Gm05_3661638, Gm06_47199506, and Gm07_7868756.

The significant markers found from this study do not overlap with the signals found for grain yield (Diers et al. 2018) and yield stability (Xavier et al. 2018) in this population. However, markers Gm02_639640, Gm07_7868756 and Gm12_2838455 are near seed size QTLs reported by Diers et al. (2018). Two markers, Gm19_1587494 and Gm18_57370051, were found to be associated with important traits from previous studies.

The marker Gm19_1587494 was also found to be the key association to canopy coverage (Xavier et al. 2017b), which means that canopy coverage could be associated with the three yield components. The marker Gm18_57370051 is linked to the stem termination gene Dt2 (Bernard et al. 1972), which has been previously detected in NAM families by Ping et al. (2014). In previous studies, Hao et al. (2011) and Fang et al. (2017) found that Dt2 is an influential gene on the number of pods and nodes. The Dt2 gene is also believed to have played a role in the soybean domestication (Sedivy et al. 2017).

The markers that were found to be associated to yield components in this study had little to no impact on maturity, which can be a major limiting factor to their use in breeding as most QTLs that improve yield often increase the number of days to maturity (Table 2). However, the QTL peaks also had a limited impact on grain yield across families, with effects ranging from −0.46 to 0.62 bu/ac.

### Genomic selection

Markers are informative at two levels for genomic predictions: they can inform the relationship and detect markers linked to, or in linkage disequilibrium with, the quantitative trait loci (Habier et al. 2007). Within-family predictive ability solely relies on the linkage between markers and QTLs, as the relationship among individuals is constant. The predictions of families not included in the training set (leave-family-out) can yield mixed results since the training set often holds families with shared ancestry. Of course, the controlled ancestry is a key property of NAM populations since all families share a common parent and, therefore, the outcome predictive ability is higher than the non-experimental population where neither parent has offspring in the calibration set. Predictions performed across family are presumably the most likely to be accurate, as they capture relationships among families and disequilibrium between markers and QTLs.

Figure 3 depicts well the expected predictive ability, as within-family predictions hold a high degree of uncertainty, with correlations averaging from 0.064 across yield components, followed by leave-family-out predictions, with an average correlation of 0.092, and the most predictive was across-family predictions, with average correlations above 0.224. Note that prediction across families can also be misleading as these are subject to the Simpson paradox (Chipman and Braun 2017), where the model is able to detect large differences across families that inflate the total correlation, but the predicted families may display negative correlation within-family.

The difference in predictive ability between GBLUP and BayesB, which translated into an average improvement of 0.02 going from GBLUP to BayesB, is due the larger flexibility the BayesB model, which is more likely to capture large and null effects (Meuwissen et al. 2001, Habier et al. 2011, Perez and de los Campos 2014, Xavier et al. 2016). Having a comparison between GBLUP and BayesB can provide an insight into the genetic architecture of the trait under evaluation (Daetwyler et al. 2013). In this study, we expected BayesB to outperform GBLUP since we uncovered a partially oligogenic architecture from the association analysis, but a key piece of information that the genomic prediction analysis provides is discrepancy between GBLUP and BayesB, which inform the degree to which the genetic architecture of the traits under evaluation depart from a polygenic architecture.

Note that the advantage provided by changing the model from GBLUP to BayesB is not comparable to the difference in predictability between cross-validation methods (i.e. within-family, leave-family-out, and across-family). The reason why this phenomenon occurs is that different methods may improve how well the model detects the genetic architecture, but different types of cross-validation provide different information. Thus, gains associated to the choice of a prior are often considered negligible in comparison to increases in population size, better experimental practices, or more representative calibration sets (de los Campos et al. 2013, Xavier et al. 2016). A possible way of capturing more information for genomic prediction is the explicit modeling of other sources of genetic information, such as dominance and epistasis (Xu 2013). As presented in this study, yield components in some populations have a greater influence of epistasis than the additive background and, on average, the within-family variance decomposition indicates that additive genetics explains as much of the yield components phenotypes as epistasis (Figure 4).

### Stability and plasticity

The genetic correlations across-environments within-trait (e.g. pods 2013 and pods 2014) were larger than the “additive” counterparts (Figure 5B), ranging from approximately 0 to 0.4, whereas the additive correlations ranged from 0 to 0.25. The discrepancy between genetic and additive genetic correlations is attributed to the genetic control due to QTLs and non-additive polygenic genetic background.

For the families with near-zero genotype-by-environment correlation, performing selections with a single year of data may not translate into observable genetic gains in the subsequent years, and collecting data from more environments may not necessarily increase the predictive ability of the yield components. Particularly for yield components, low genotype-by-environment correlations is not necessarily bad since soybean yield plasticity relies on distributing resources among yield components, which serves as a physiological response to mitigate yield losses under stress (Board and Tan 1995, Board et al. 1997, Pedersen and Lauer 2004, Zhang et al. 2004). Whereas yield components are mainly responsible for the yield formation, these are not necessarily the best linear yield predictors (Board and Modali, 2005). For example, Board and Harville (1993) showed that the number of pods serves as the mechanism by which seed production increases in response to greater light interception.

Our previous study (Xavier et al. 2017a) assessed the association among soybean agronomic traits and yield components in the SoyNAM population based on undirected graphical models. The graphical models depicted genetic and environmental interdependence among yield components. That means that interactions among yield components occur due to genetic forces as well as a response to environmental stimuli and agronomic practices. Such a phenomenon is also described in a summary of agronomic studies on soybean yield components authored by Board and Kahlon (2011). The interactions among yield components play a key role in the redistribution of resources and yield stability (Ball et al. 2000). It is possible that breeding any given yield component towards extreme values may result in a compromised ability of soybeans to compensate yield under stress (Malausa et al. 2005).

### Yield increases

From the standpoint of trait decomposition, the breaking down of grain yield into pods and nodes does not seem to be an effective approach since there is no strong evidence that these yield components are more heritable than yield (Table 1) or strong genetic correlation to yield (Figure 5A) that would justify the selection based on yield components. Except for a few families, yield components are not good proxies for grain yield (Figure 5C). It is possible that the genetic architecture of the yield components under evaluation is just as complex as grain yield itself, not justifying predicting yield components instead of yield *per se*.

In our previous genomic prediction study (Xavier et al. 2016), we assessed how a variety of different genomic prediction models would predict the agronomic traits and yield components under the following scenario: within year and across-population. Even though that study did not provide in-depth insight into the genetic architecture of yield components, it was found that genomic prediction models that can jointly account for large effect QTLs and epistasis were advantageous over simpler prediction approaches. That study also found that predicting yield is easier than predicting yield components. Those results were further confirmed by the current study, where we assessed the architecture of yield components with more data and under different approaches.

### Phenotyping

A major challenge of working with yield components is the data collection as the counting is highly subjective to human error, lowering the trait heritability and affecting the signal detection in downstream analysis. As deep learning methods for computer vision become increasingly population for phenotyping morphological traits (Singh et al. 2018), the current limitations with data collection could be addressed by an automated high-throughput phenotyping instead of human counts, that would likely increase both accuracy and scalability of the process. For instance, both Uzal et al. (2018) and Li et al. (2019) recently proposed an imagery system for counting seeds directly from images of soybean pods, yet another yield component limited by the challenging phenotyping. Technologies that enable better, faster and cheaper data collection remain a key limiting factor to research on yield components.

## Authors contribution statement

AX and KMR planned the experiment. AX collected and analyzed the data and wrote the manuscript. KM provided insight on the data analysis and review the manuscript.

## Acknowledgments

We thank the SoyNAM collaborators for their contributions to the experiment. William Beavis for experimental design, Qijan Song and Perry Cregan for genotyping, and Jim Specht and Brian Diers for creating the germplasm resource. We thank Chris Hoagland and Curtis Brackett for managing the experiments and contributed to collect the phenotypes. We thank the United Soybean Board for funding the field experiment in 2013 and Corteva AgriSciences for funding the data collection in 2013 and 2014, and the field experiment in 2015.

